# The unbalanced distribution of the amino aicds(Arg, Lys) in high pI proteins of complete proteomes, its origin and evolution

**DOI:** 10.1101/090290

**Authors:** Zhenhua Xie

**Affiliations:** The Shenzhen Key Laboratory of Health Sciences and Technology, Graduate School at Shenzhen, Tsinghua University, Shenzhen 518055, China.; Green biosynthesis institute of Bontac bio-engineering(shenzhen)Co.,Ltd, Shenzhen 518102, China

## Abstract

No evolutionary signature has been found in the complete proteomes of eukaryotes by now, although amino acid composition signature of the complete proteomes was discovered as molecular signature of the adaptation for thermophiles and halophiles. Arginine and lysine respectively have the guanidinium group and the ∊-amino group as the ionizable side chain groups with different pKa values of about 12.5 and 10.5. The trends of their distribution seem similar in the range of about pIs < 10.0 and diverge in the range of about pIs≥10.0 in most complete proteomes of 387 species from the three domains of life. The complete proteome of Reticulomyxa filose is the one of only in 287 eukaryotic complete proteomes that has a predominance of the trend of lysine over that of arginine in high pI proteins. The unbalanced distribution of the amino aicds(Arg, Lys) in high pI proteins of complete proteomes may originally come from different pKa values of arginine and lysine and be developed by the influences of an average lysine level of a proteome and evolution. Because of this unbalanced distribution, the pattern of arginine and lysine distribution in high pI proteins of some complete proteomes can form a particular proteomic structure as evolutionary signature that may be shaped by massive natural selection in molecular level from hundreds to ten thousands of proteins in the complete proteomes of many animals and green plants.

---

A complete proteome is a unit of proteins expressed by a genome completely sequenced. Sequences of the proteins in a complete proteome can be translated from all protein coding genes of a genome completely sequenced^1, 2^. As essential parts of organisms, proteins have many functions and participate in every process within living organisms, for example, catalyzing the metabolism of proteins, RNA, DNA and metabolic substances, transducing biological signals, transporting biological molecules^3, 4^. The availability of increasing numbers of complete proteomes provide a precious resource for deciphering and interpreting basic mechanisms of molecular evolution on a large scale in life on Earth.

Many studies have identified amino acid comopsitions of extremophiles related to their environmental niches, and some results have revealed that the amino acid compositions of the adaptive complete proteomes are associated with the evolutionary adaptation of extreme environments, for example, high temperature and high salinity ^5–8^. Therefore, retainment of protein structural and functional integrity may be essential for extremophiles to survive in extreme environments.

All above-mentioned studies have used average value of the amino acid compositions of complete proteomes to explore the proteome evolution and lifestyle adaptation. Until now, any trend of amino acid composition distribution has never be used to to explore the proteome evolution. It has be exciting that the unique trend of arginine and lysine distribution versus the protein pI values of complete proteomes was discovered.

The isoelectric point (pI) of a protein can be defined as the pH value of solution at which its total electrical charge is zero^3^. A protein can be precipitated from the solution by adjusting the solution pH to the protein’s isoelectric point because of protein’s isoelectric aggregation. Proteomic analysis has extensively used 2-D gel to separate proteins according to molecular weight(MW) and pI’s. Protein pIs can be experimentally measured by polyacrylamide gel-based isoelectric focusing. Based on the assumption that the protein would be denatured and the pKa values of ionizable amino acid residues would be constant irrespective of structural context, theoretical protein pIs can be precisely calculated from the sequence of a protein. The results of 2-D gel have showed that theoretical pI values match very closely to the protein pIs measured experimentally^9–12^.

The isolectric point(pI) of a protein depends only on C-terminal carboxylic acid group, N-terminal amino group and ionizable side-chain acid-base groups that can be free to ionize. The amino aicds (Asp,Glu,Arg,Lys,His, Cys,Tyr) with an ionizable side-chain have an additonal acid-base group^3^. Compared with the low-abundant amino acids(His, Cys, Tyr), amino acids(Asp,Glu) or (Arg,Lys) are similar with each other in some physicochemical properties and high abundant in complete proteomes^3, 13, 14^. Therefore, amino acids(Asp,Glu,Arg,Lys) have more evolutionary potential beyond maintaining protein structure and function integrity. In this paper, arginine (Arg) and lysine(Lys) have be chosen to analyze the distribution of the their abundance values versus the corresponding values of theoretical pIs in 387 complete proteomes.

## Methods

All complete proteome sets in FAST format for 387 species were downloaded from the Universal Protein Resource (UniProt) in early 2015. Using the R statistical programming language, the original sequences of proteins in FAST format were converted into the M-truncated sequences(MTSs) that are in plain-text format without the initial methionine, and the values of abundances of amino acids (Arg, Lys) in the MTSs were calculated. The values of pIs of all MTSs were computed using Compute pI/Mw tool (http://web.expasy.org/compute_pi/). Based on the values of theoretical pIs and abundances of amino acids (Arg,Lys), the map of the distribution of one amino acid (Arg or Lys) for pIs in any complete proteome had been illustrated by the R statistical programming language. In order to clearly display the trend of an individual amino acid distribution, the red average line and green loess curve had been introduced into the maps. The loess curve can be simply used as a trendline to demonstrates the trend of a scatterplot.

## Results

In this study, the maps of arginine and lysine distribution versus pIs in complete proteomes of 387 species from the three domains of life were constructed and all demonstrated in three additional files in which the maps in complete proteomes from archaea(50 species), bacteria (50 species) and eukaryotes (287 species) list respectively. The additional files were named as “maps in eukaryotes”, “maps in bacteria” and “maps in archaea”. Here, the maps in Figure 1 and Figure 2 were selected from the additional files to illustrate the discoveries in this paper.

**Figure 1.**
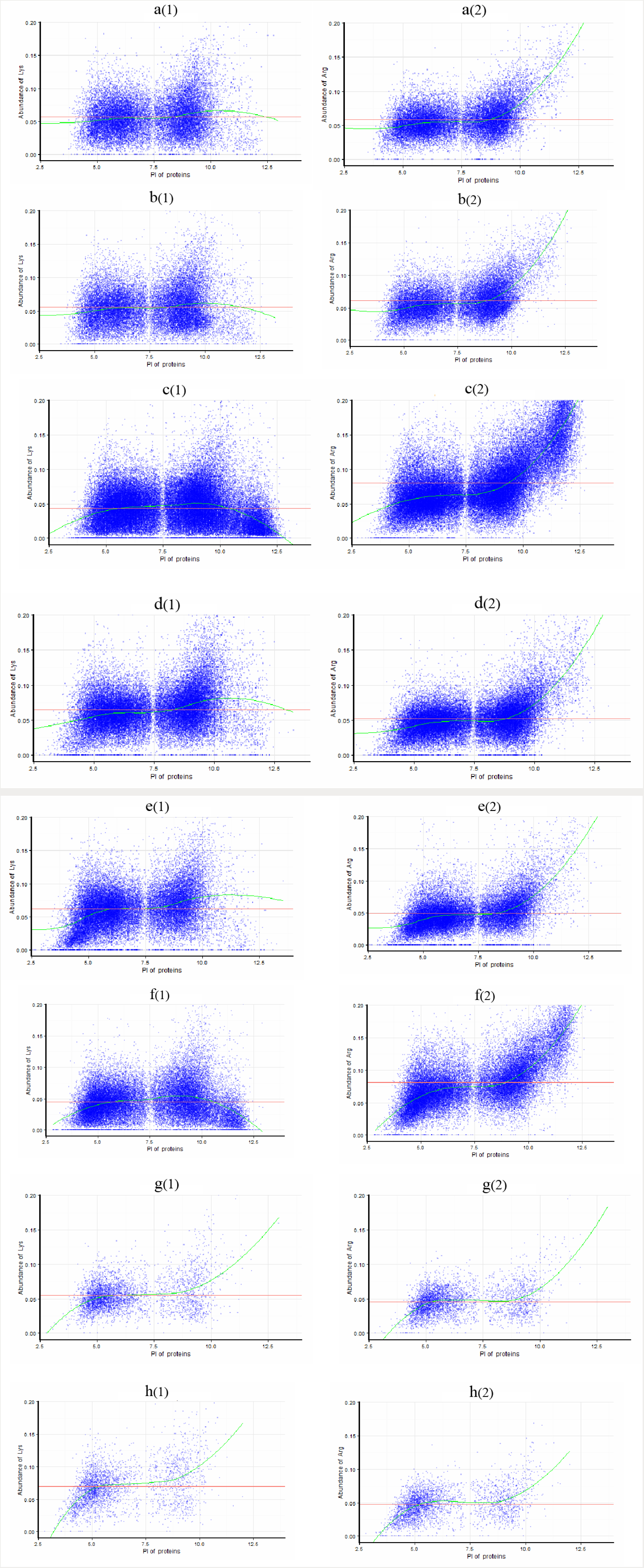
The maps of the distribution of Arg and Lys versus pIs for the eight complete proteomes. (a) chimpanzees (Pan troglodytes); (b) platypus(Ornithorhynchus anatinus); (c) asian rice (Oryza sativa subsp. japonica); (d) tomato (Solanum lycopersicum); (e) sponge (Amphimedon queenslandica); (f). marine diatom(Thalassiosira oceanica); (g) Vibrio sinaloensis; (h) Methanosarcina sp. MTP4.

**Figure 2.**
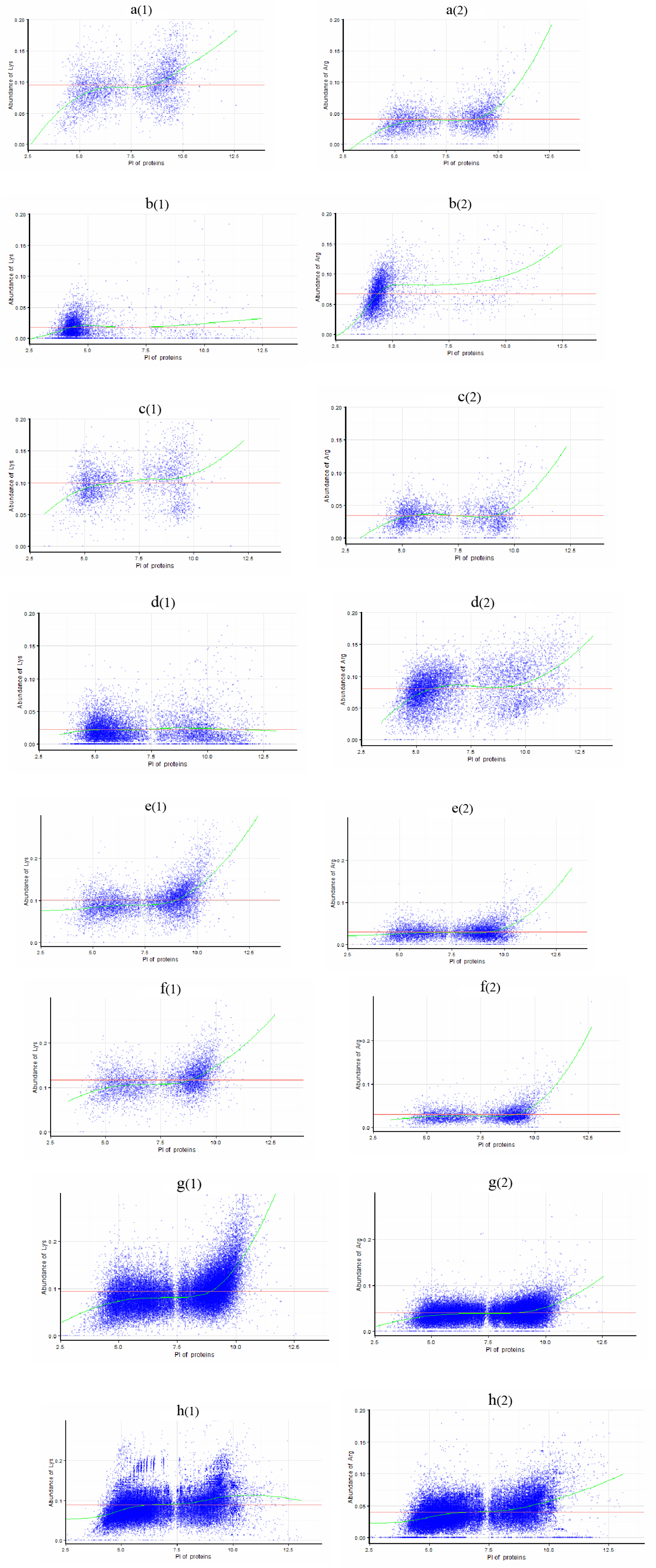
The maps of the distribution of Arg and Lys versus pIs for the eight Lys-rich and Lys-low complete proteomes. (a) archaeon Loki; (b) Haloterrigena turkmenica; (c) Clostridium baratii str. Sullivan; (d) Amycolatopsis mediterranei; (e) Ichthyophthirius multifiliis; (d) Plasmodium falciparum; (e) Reticulomyxa filose; (f) Trichomonas vaginalis.

### The unbalanced distribution of the amino aicds(Arg, Lys) in high pI proteins of most complete proteomes

By the comparison between arginine and lysine distribution versus pIs in these maps of 387 complete proteomes, an obvious feature has been discovered that the trends of arginine distribution are similar with that of lysine distribution except in a range of high pIs(about pIs≥10.0) and different from that of lysine distribution in most complete proteomes in a range of high pIs(about pIs≥10.0). This different distribution of the amino aicds(Arg, Lys) in high pI proteins embedded deeply in most complete proteomes has been designated as the unbalanced distribution (see the Figure 1).

In fact, the predominance of the trend of arginine over that of lysine in high pI proteins exists in most complete proteomes, chimpanzees (Pan troglodytes), platypus(Ornithorhynchus anatinus), asian rice (Oryza sativa subsp. japonica), tomato (Solanum lycopersicum), sponge (Amphimedon queenslandica) and marine diatom(Thalassiosira oceanica), for example(see the Figure 1). Only a few complete proteomes have a good concordance between the trends of their distribution in the all range of pIs, Vibrio sinaloensis and Methanosarcina sp. MTP4, for example(see the Figure 1).

### The unbalanced distribution might be affected by an average lysine level of a proteome

Although the complete proteomes of archaeon Loki, Trichomonas vaginalis, Haloterrigena turkmenica and Amycolatopsis mediterranei have different average lysine levels of abundance 0.0953, 0.0945, 0.0180and 0.0224, the trends of arginine are respectively more steep than that of lysine in high pI proteins of these four complete proteomes. Because of their low average lysine level, it is hard to maintain a good concordance between the trends of arginine and lysine distribution except in a range of high pIs(about pIs≥10.0) in the complete proteomes of Haloterrigena turkmenica and amycolatopsis mediterranei (see the Figure 2).

Because of their high average lysine level of abundance 0.0993, 0.1007 and 0.1168, a good concordance between the trends of arginine and lysine distribution can be found in all range of pIs in the complete proteomes of Clostridium baratii str. Sullivan, Ichthyophthirius multifiliis and Plasmodium falciparum (see the Figure 2). The complete proteome of Reticulomyxa filose in an average lysine level of abundance=0.0945 is the only one of 287 eukaryotic complete proteomes that has a predominance of the trend of lysine over that of arginine in high pI proteins (see the Figure 2). Therefore, average lysine level of a proteome might somehow affect the unbalanced distribution.

### The origin of the unbalanced distribution

In order to find the origin of the unbalanced distribution, the five complete proteomes of Archaeon Loki, Amycolatopsis mediterranei, Thalassiosira oceanica, Zootermopsis nevadensis and Pan troglodytes were choose as imitated proteomes to create their imitating random-proteomes, and the five imitating random-proteomes were contructed by the way that the proteins in every slice of a imitated proteome were replaced by representative random protein sequences in identical number. The five imitating random-proteomes were designated as random-proteome(Loki), random-proteome(AM), random-proteome(TO), random-proteome(ZN) and random-proteome(pan).

The minor predominance of the trend of arginine over that of lysine in high pI proteins exists in the five imitating random-proteomes, so the unbalanced distribution does clearly exist in the five imitating random-proteomes (see the Figure 3). This result has suggested that the origin of the unbalanced distribution may be traced to different pKa values of arginine and lysine that could affect the unbalanced distribution to such a degree that the trend of lysine distribution in a range of high pIs is similar with that of lysine distribution and still rises.

**Figure 3.**
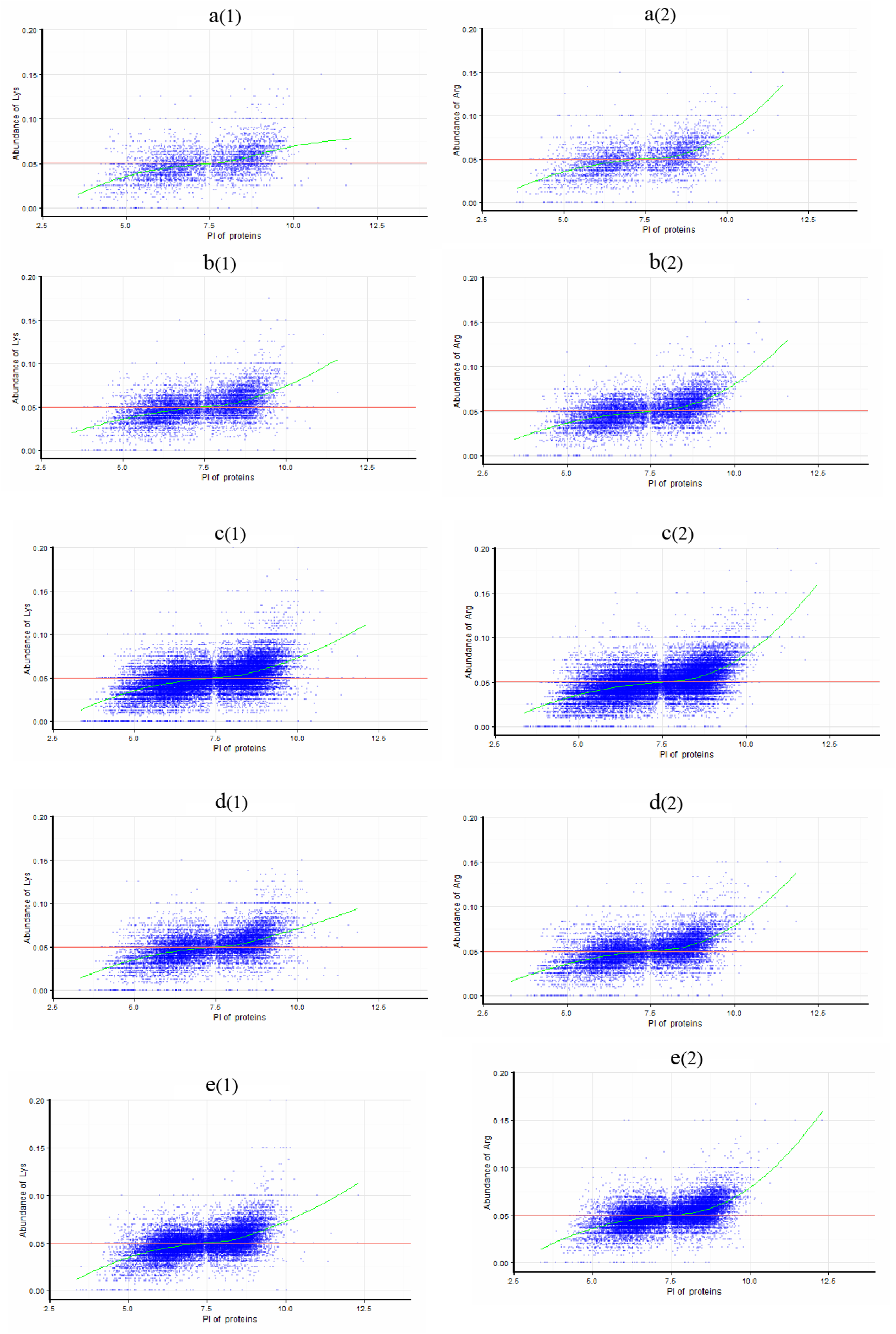
The maps of the distribution of Arg and Lys versus pIs for the five random-proteomes. (a) random-proteome(Loki) (4061 random sequences); (b) random-proteome(AM) (9551 random sequences); (c) random-proteome(TO) (34547 random sequences); (d) random-proteome(ZN) (14539 random sequences); (e) random-proteome(pan) (20131 random sequences).

### The evolution of the unbalanced distribution

For exploring potential evolutionary trend of the unbalanced distribution, a complete proteome is defined as typeⅠif the trend of lysine distribution in a range of high pIs rises and as typeⅡ if the trend of lysine distribution extends horizontally, downward or even goes up first and then down. Based on this definition, the five imitating random-proteomes belong to typeⅠ,and the unbalanced distribution in a typeⅡcomplete proteome could be shaped by an evolutionary force.

There are 44 typeⅠand 6 typeⅡcomplete proteomes in 50 archaea, 40 typeⅠand 10 typeⅡ complete proteomes in 50 bacteria and 183 typeⅠand 104 typeⅡcomplete proteomes in 287 eukaryotes. In archaea, bacteria and eukaryotes, the proportion of typeⅡcomplete proteomes were respectively accounted for about 12%, 20% and 36.2%.

## Discussion and Conclusion

In a complete proteome, high isoelectric point proteins may play a critical role in molecular interactions with negative-charged molecules, for example, DNAs, RNAs, negative-charged proteins and small negative-charged molecules. The distributions of strongly basic residues(Arg,Lys) in high isoelectric point proteins of complete proteomes may provide new insight into proteome evolution.

In high pI proteins of the most complete proteomes, the trend of arginine distribution rises while the trend of lysine distribution rises gradually, extends horizontally or downward and even goes up first and then down. This difference between arginine and lysine distribution in high pI proteins may be concerned about a distinction between arginine and lysine function in life.

In a typeⅠcomplete proteome, lysine has a similar upward trend to that of arginine. The five imitating random-proteomes belong to typeⅠ. The minor predominance of the trend of arginine over that of lysine in high pI proteins of the five imitating random-proteomes demonstrate that the different pKa values of arginine and lysine could slightly affect the unbalanced distribution. In a typeⅡcomplete proteome, the trend of lysine distribution is not similar to that of arginine. The trend of lysine distribution of a typeⅡcomplete proteome is so different that the effect of the different pKa values of arginine and lysine can not reach to such an extent. Therefore, a force has been involved in the formation of the unbalanced distribution in a typeⅡcomplete proteome.

In archaea, bacteria and eukaryotes, the proportion of typeⅡcomplete proteomes were respectively accounted for about 12%, 20% and 36.2%. Therefore, the unbalanced distribution in a typeⅡcomplete proteome may reflect the result of evolution of this species.

In the range of high pIs(about pIs≥10.0) of the complete proteomes of many animals and green plants, the centralized distribution of abundant arginine and relatively decentralized distribution of low abundant lysine form a particular proteomic structure. This particular proteomic structure has been shaped by massive natural selection in molecular level of from hundreds to ten thousands of proteins in the complete proteomes of many animals and green plants. The pattern of arginine and lysine distribution in the range of about pIs≥10.0 in the complete proteomes of asian rice (Oryza sativa subsp. japonica) and marine diatom(Thalassiosira oceanica) can clearly illustrate this particular proteomic structure. What is evolutionary force to drive this massive natural selection between the strongly basic residues(Arg,Lys)?

In the complete proteomes of chimpanzees, platypus, asian rice, tomato, sponge and marine diatom, the number of high pI proteins is 937, 1650, 12236, 2151, 1237 and 5618 respectively. It is hard to design a experiment to search some evolutionary force to drive this massive natural selection between the strongly basic residues(Arg,Lys).

In this paper, a hypothesis has been put forward to explain the cause of the unbalanced distribution. The difference between arginine and lysine distribution in a range of high pIs may be involved with the distinction between arginine and lysine posttranslational modification in proteins. The pKa of the ∊-amino group of lysine is lower than that of the guanidinium group of arginine, so lysine is a better nucleophile and more favorable than arginine for posttranslational modifications in proteins. For example, ubiquitination and acetylation were identified only as posttranslational modifications of lysine. The ubiquitination and acetylation/deacetylation can respectively cause protein degradation via the proteasome and energy consumption^15, 16^. Low level of the ubiquitination and acetylation/deacetylation may be required for protein mechanism and cell integrity. High level of the ubiquitination and acetylation/deacetylation may unproperly cause promotion of protein unstability and the futile cycle. Therefore, the predominance of the trend of arginine over that of lysine in a range of high pIs(about pIs≥10.0) in most complete proteomes may be beneficial for life to survive.

Now that the predominance of the trend of arginine over that of lysine in a range of high pIs(about pIs≥10.0) may has a benefit for life to survive, it is hard to explain why is the trend of arginine distribution similar with that of lysine distribution except in a range of high pIs(about pIs≥10.0). There might exist a enzymatic system that is specific to catalyze the ubiquitination and acetylation of the proteins in a range of high pIs(about pIs≥10.0).

It is necessary to explore what evolutionary force and mechanism that have caused the unbalanced distribution of arginine and lysine in complete proteomes.

## Acknowledgements

The author would like to acknowledge support from Shenzhen Bureau of Science,Technology and Information (Grant No. JCYJ20140417115840267 and JCYJ20150518162154828).

## Author Contributions

Zhenhua Xie did all works in this study and wrote the manuscript.

